# Secreting salt glands constrain cuticle fracture to enhance desalination efficiency

**DOI:** 10.1101/2025.02.27.640653

**Authors:** Melissa H. Mai, Fulton E. Rockwell, Juan M. Losada, Nya Nicholson, Zhigang Suo, N. Michele Holbrook

## Abstract

Plants responding to excessive soil salinity by discharging brine onto their leaf surface risk dehydration through the osmotic continuity between the living tissue and the surface brine, which further enriches with evaporation. Cuticle cracks have long been identified as essential for salt to reach the leaf surface but provide the potentially desiccating continuity between the brine and the gland interior. Using the secreting salt gland of *Nolana mollis* as a model system, we integrate mathematical modeling, imaging, and physiological measurements to examine the mechanical and biochemical processes required for efficient desalination. We find that the subcuticular space between the concentrated surface brine and the more dilute secreting cell eases the energetic limits of active desalination by reducing the concentration gradient of salt across the cell membrane. We show that crack size plays a critical role in balancing the osmotic and pressure gradients required for salt removal without runaway foliar desiccation.

---

Soil salinity poses both a biochemical and a physical challenge for terrestrial plants. Elevated exposure to salt interferes with proper osmoregulation and induces protein malfunction and aggregation, leading to stunted growth, impaired development, and increased mortality [1–6]. As ions are transported through the xylem, higher rates of transpiration exacerbate salt accumulation within the plant [7]. Yet, preventing this accumulation by exclusion at the roots produces large osmotic gradients that can impair hydraulic function [8]. To address the full range of issues presented by salt stress, strategies for salt tolerance are expected to encompass not only biochemical but also physical solutions.

Halophytic plants have evolved specialized mechanisms for salt tolerance and exist within several plant lineages [9, 10]. Many halophytes address belowground salt stress by excluding salt from their roots or by accumulating salt in the roots to reduce accumulation in the leaves [11–14]. However, for most plants, there is often not enough water flow around the roots to wash away the buildup of excluded salts. As salts accumulate in the root zone, replacement of fouled roots incurs significant energetic costs for the plant [6]. Notable exceptions are mangroves, whose roots grow in waterlogged soils where mobile seawater can advectively or diffusively dilute the locally hypersaline water [12, 15]. Nevertheless, salt absorption cannot be fully avoided, and accumulation of excess salts leads to the induction of leaf succulence until the salts are physically cleared from the plant during leaf senescence [15].

Recretohalophytes rely less on exclusion and instead use energy to drive elimination of salt from their leaves via specialized cellular structures known as salt glands. Present in many plant genera, salt glands are morphologically and functionally diverse, with two main strategies for salt removal [16]. One strategy is sequestering salt from other tissues in the vacuoles of bladder-like cells that are eventually shed [17]. The other strategy is secretion of excess salt onto the leaf surface (exo-recretohalophytes) [18]. Secretory cells at the gland apex actively deposit salt beneath the cuticle, either through membrane transporters or exocytosis of microvacuoles [19, 20]. The resulting osmotic gradient across the cell membrane pulls water from the gland into the subcuticular space, generating pressure [21]. Cracks in the cuticle allow the salt solution to reach the leaf surface, where it becomes further concentrated by evaporation into a brine [22–24]. These cracks also form an aqueous continuum between the resulting surface brine and the living tissue. How these plants manage to prevent self-desiccation despite this strong osmotic gradient is not understood.

Here we examine how the secreting salt gland’s multicompartment morphology contributes to the system’s success. We develop a steady-state mathematical model of the secreting salt gland to determine the role of the intermediate subcuticular space in mediating salt removal without catastrophic water loss. We find that the success of secreting salt glands relies not solely on biochemical processes operating at the cell membrane but also on their structural organization and the fracture mechanics of the cuticle.

## RESULTS

### Ultrastructure of the secreting salt gland

We use *Nolana mollis* (Phil) Johnston as the model species for our analysis of secreting salt glands. *N. mollis* is a succulent-leaved shrub dominating the Pan de Azúcar coastal valley of the Atacama Desert (Fig. 1a) and is distinctively covered with a persistent and concentrated brine (Fig. 1b) while neighboring species remain dry [25, 26]. *N. mollis*’s deep taproot provides access to a consistent, but saline, source of groundwater [27]. Secretion of the brine maintains a stable whole-leaf osmolality (500 mmol/kg) despite high soil salinity (Fig. S1).

**Fig. 1.**
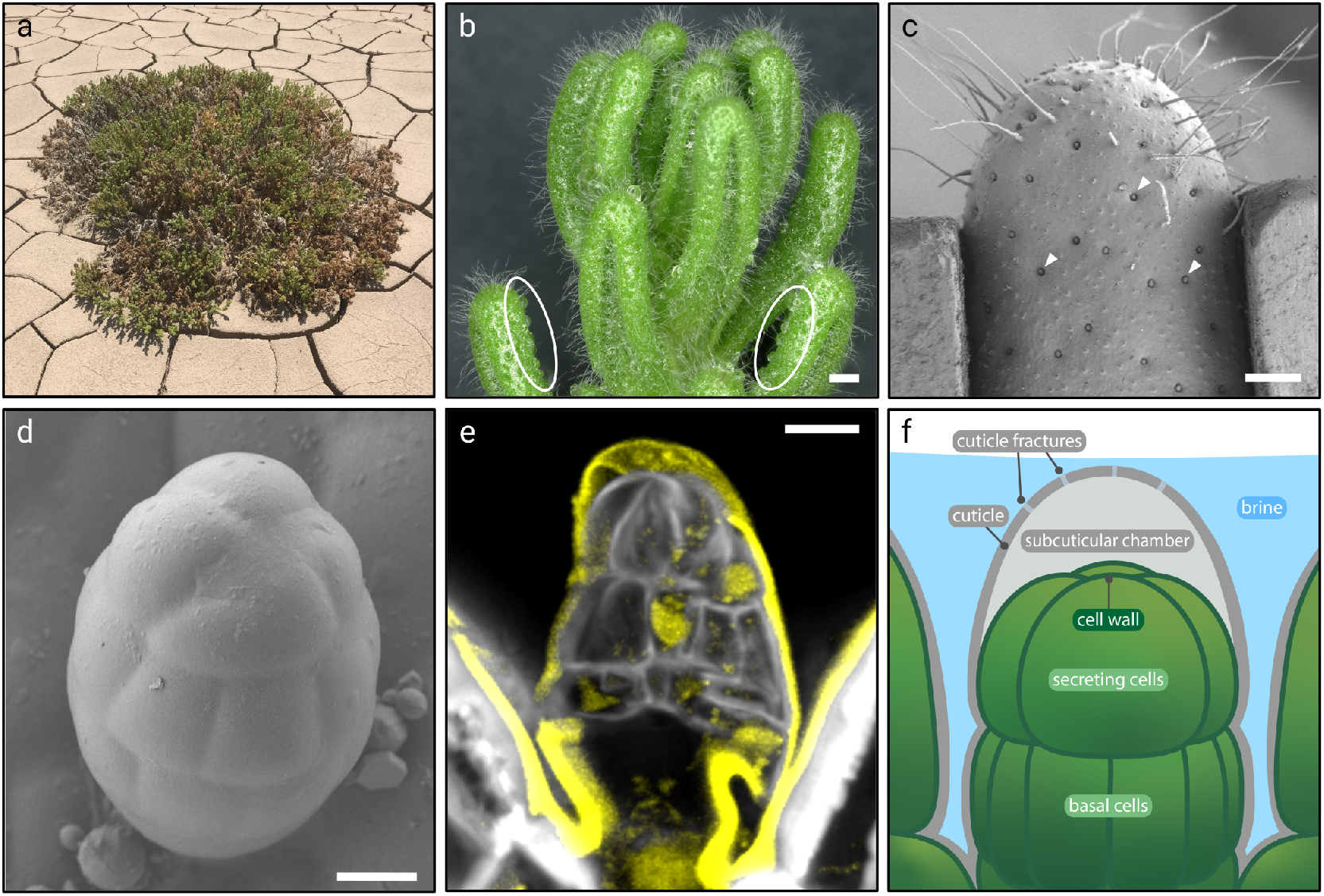
Salt gland structure of *Nolana mollis*. a, *N. mollis* is a shrub native to Pan de Azúcar, Atacama Desert, Chile. b, The leaves of *N. mollis* secrete brine onto their surface (circled). c-d, Cryo-SEM imaging of the adaxial leaf surface (c) reveals multicellular brine-secreting salt glands (d) in depressions of the epidermis (c, white arrows). e, Confocal imaging of the salt gland highlights a chamber formed between the cuticle (yellow) and the cell wall (white) at the gland apex. f, Conceptual diagram of the chambered secreting salt gland ultrastructure. Cracks in the cuticle allow for aqueous salt to be secreted into the surface brine. Scale bars: 1 mm (b), 400 *µ*m (c), 10 *µ*m (d,e).

The salt glands, with a diameter of 20-30 *µ*m, lie in depressions of the leaf epidermis and cover approximately 1% of the leaf surface (Fig. 1c,d). Confocal imaging of the salt gland (Fig. 1e) reveals the cuticle (yellow) visibly detached from the cell wall (white), forming an inflated subcuticular chamber at the gland apex. Details of the ultrastructure of the *N. mollis* salt gland are summarized in Fig. 1f.

The conceptual framework of the secreting salt gland is presented in Fig. 2. Salt moves from the secretory cells into the chamber via active processes. The details of this remain unclear, as specific ion transporters involved in foliar salt glands have yet to be identified [28–30]. Microvesicular transport has also been hypothesized but not yet confirmed [31]. Regardless of the specific mechanism, movement against an ion gradient requires an input of energy [32], whether to induce protein conformational changes [33], operate proton pumps [34–36], or drive vesicular formation and movement [31]. Histological studies have consistently found mitochondrial enrichment in secretory cells, revealing the substantial energetic cost of salt removal [19, 20, 23, 24, 37, 38]. When the concentration gradient across the cell membrane becomes steep enough, the energetic cost of moving an ion against its gradient becomes too large, and export activity vanishes. A theoretical response of transporter activity to the concentration gradient across the membrane is sketched in Fig. 2c. The exact shape of this curve is unknown for real systems, and we find that, as long as the response vanishes at a large concentration gradient, our results are qualitatively unaffected (Fig. S2).

**Fig. 2.**
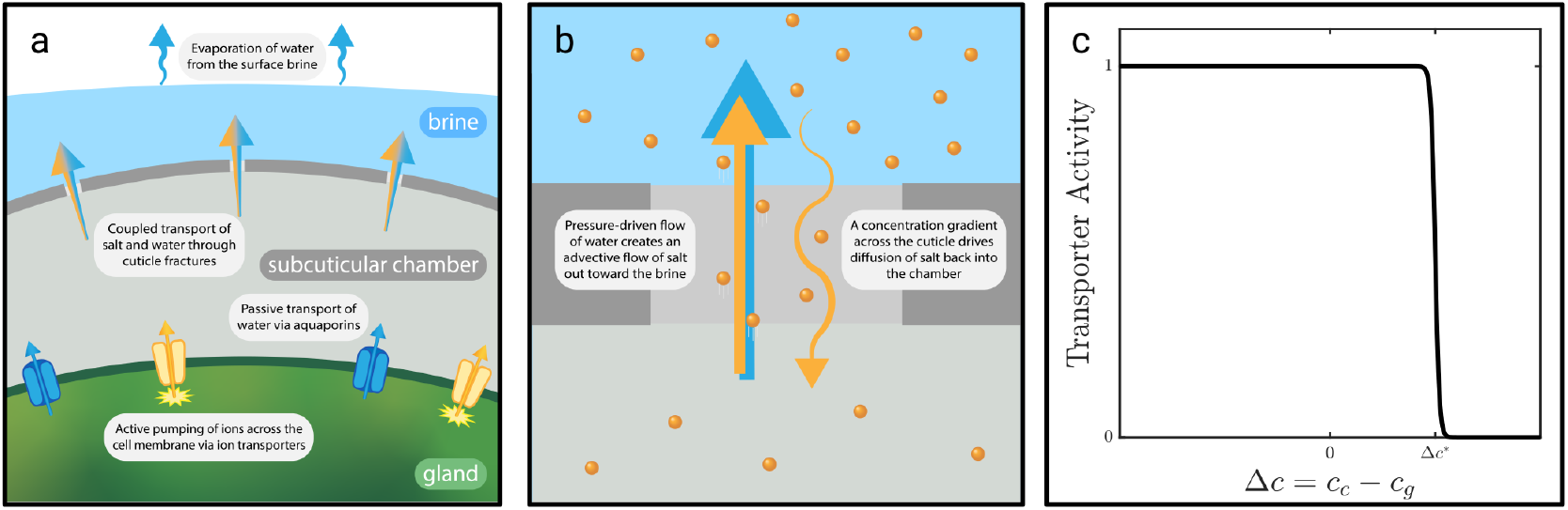
Conceptual framework for the secreting salt gland model. a, Salt (orange) is actively loaded into the chamber while water (blue) moves passively across the cell membrane. The aqueous salt solution in the chamber reaches the surface through cracks in the cuticle. Evaporation from the leaf surface concentrates the surface solution into a brine. b, Through cracks in the cuticle, pressure-driven flow of water can move salt advectively out toward the brine (straight arrows). A concentration gradient across the cuticle drives diffusion of ions through the cracks (curved arrow). If the brine is more concentrated than the chamber, diffusion will run counter to the advective flux. c, Theoretical response of ion transporter activity in the cell membrane as a response to a concentration gradient Δ*c* across the membrane between the gland and the chamber. Transporter activity vanishes above a threshold value Δ*c*^*^.

The electroosmotic gradient established across the cell membrane by ion transporters drives an osmotic flow of water into the chamber via membrane-bound aquaporins. The resulting buildup of pressure inside the chamber drives advective flow of the enclosed salt solution out toward the exterior brine via cracks in the cuticle. Diffusion of salt also occurs through these channels. If the concentration of the brine exceeds that of the chamber, the net diffusive flux will drive ions back into the chamber, counter to the advective current (Fig. 2b).

The volume and concentration of the surface brine is modulated by evaporation, crystallization of salts, and dripping of the brine off the leaf surface [25]. Pre-dawn brine osmolality was measured to be around 2000 mmol/kg (Fig. S1). We expect the concentration gradient between the brine and the gland, if not separated by the intermediate chamber, to be too steep to be overcome by active transport processes. In our model, we consider the steady state, with the brine concentration set to its empirical value, to determine the contribution of the intervening subcuticular space to the efficiency of salt expulsion as a function of cuticle crack size.

### Model results

Model results are illustrated in Fig. 3. When cracks are small, pressure builds up in the chamber, and when cracks are too large to contribute meaningful resistance to flow, the pressure relaxes to atmospheric levels (0.1 MPa) (Fig. 3a). The total water flux, corresponding to the water lost from the gland, increases with larger crack sizes, eventually plateauing as the increasing crack size corresponds to the complete destruction of the cuticle (Fig. 3b). Considering salt gland coverage at approximately 1% of the leaf surface, the maximal water loss from the salt gland is about 0.3 mL/cm^2^/day, or 3 mm/day, comparable to the potential evapotranspiration of arid lands (4-6 mm/day) [39]. Whereas transpiration is limited to times the stomata are open, gland-mediated water loss is continuous, potentially exceeding net daily transpiration and risking desiccation if the cracks are too large. Moreover, the brine itself is composed of gland secretions and nighttime deliquescence. Lower water loss rates at smaller cracks suggest that deliquescent water accounts for a larger volume fraction of the brine towards this limit. This decreases the water demand for desalination by subsidizing the total land evapotranspiration with water condensed from the atmosphere. At both the small and large fracture limits, salt concentration in the chamber exceeds the threshold for transporter activity, suppressing salt export. With large cracks, the chamber and brine become indistinguishable compartments, causing the chamber concentration to approach that of the brine. High resistance across the cuticle through small cracks impairs both water and salt export, leading to a retentive regime where salt builds up within the chamber. At intermediate crack sizes, the concentration within the chamber drops by around 40% (Fig. 3c). Within this range, advective fluxes move salt to the leaf surface faster than the diffusion of salt backward into the chamber.

**Fig. 3.**
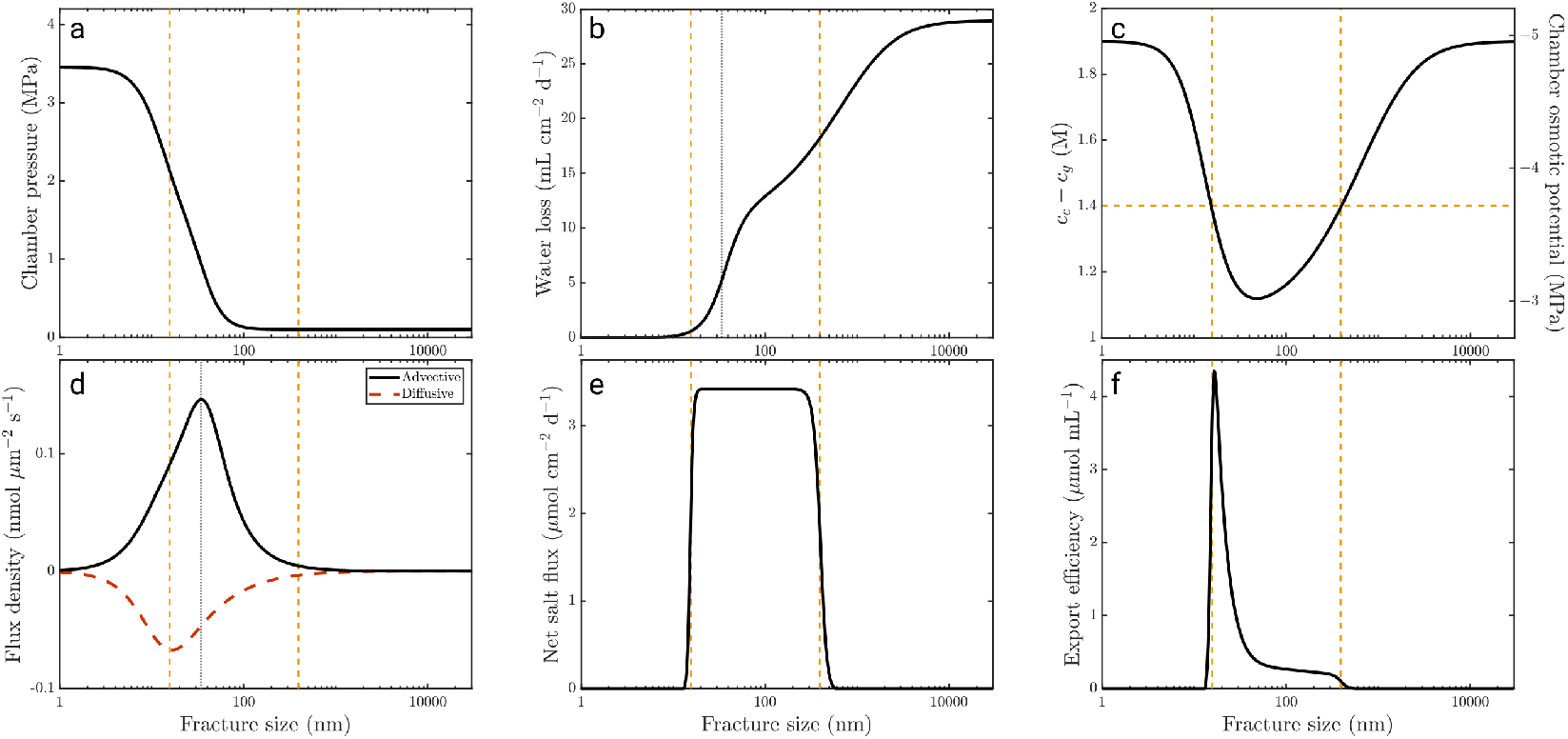
Model results as a function of fracture pore radius (*r*_*f*_, nm). a, Chamber pressure. b, Total per-area water loss from the gland, normalized by total gland area *πr*^2^ . c, Chamber salt concentration (left) and osmotic pressure (right). Ion transporters stop functioning when the concentration gradient between the chamber and the gland is above an activation threshold (horizontal orange dashed line). d, Advective (black) and diffusive (red) salt flux normalized by crack area 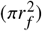,referred to as the salt flux density through the cuticle cracks. Positive flux densities indicate movement from the chamber out toward the brine. e, The net molar salt flux from the gland, normalized by total gland area, is a function of the concentration gradient across the cell membrane between the cell and chamber. f, Export efficiency (*µ*mol Na^+^ per mL water) from the gland. Vertical orange dashed lines indicate the points at which the chamber concentration crosses the transporter activation threshold. The vertical gray dotted line corresponds to the crack size with maximized advective salt flux density and the first inflection point of the water flux profile.

Fig. 3d illustrates the advective and diffusive components of the salt flux, normalized by the crack area 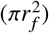,which we refer to as the flux density. While the gross advective and diffusive fluxes monotonically increase in magnitude with crack area (Fig. S3), their densities achieve local optima (Fig. 3d). Large resistance across small cracks restricts flow across the cuticle and maintains a high concentration of salt within the chamber. With increasing crack size, resistance drops, leading to larger advective fluxes, with a maximum occuring at a crack size just under 100 nm (Fig. 3d), corresponding to the first inflection point of the water flux profile (Fig. 3b). Larger fluxes more effectively clear salt from the chamber. The drop in concentration below the activity threshold (dashed line) allows ion transporter activity to resume, permitting active salt export from the secreting cell in this range (Fig. 3e). Despite increasing conductance with even larger cracks, greater crack area ultimately reduces the advective flux density by both increasing the flux area and by lowering the pressure gradient across the cuticle (Fig. 3a).

The inward diffusive flux reaches a maximum with a crack size near the lower limit of transporter activation. This corresponds to the increasing concentration gradient across the cuticle resulting from a more dilute chamber solution. With larger cracks, a smaller gradient across the cuticle, coupled with a larger crack area, reduces the diffusive flux density through the cracks. We note that the advective and diffusive flux densities scale differently with *r*_*f*_ and therefore reach their optima at different crack sizes. Within the range of crack sizes that allow for positive transporter activity, a large, outward advective flux of salt overcomes a smaller, inward diffusive flux, leading to positive salt export. Immediately outside this range, a small amount of salt export occurs, corresponding to the smooth shoulders of the chosen activity profile. Other profile shapes do not qualitatively affect the results and are explored in the SI. Beyond this range, while these flux densities are low at this limit, the large crack area allows for nonzero advective and diffusive fluxes that balance each other, resulting in negligible salt export (Fig. S3).

The export efficiency (Fig. 3f) measures the amount of salt removed (Fig. 3e) relative to the water lost from the gland (Fig. 3b). While water loss is monotonically minimized with smaller cracks, export efficiency vanishes at this limit as transporter activity halts against the steep concentration gradient. This establishes a global optimum in the export efficiency at *r*_*f*_ ≈ 20 nm, corresponding to a state in which transporters have been fully activated while the water flux remains low, one or two orders of magnitude below the potential evapotranspiration [39].

### Fracture mechanics

Cryo-SEM imaging of the salt gland cuticle revealed cracks in plants that were secreting brine (Fig. 4b-c). Glands from plants watered with dH_2_O did not secrete brine, and cracks were not observed on their cuticle surfaces. The observed crack width from active glands ranged between 10 and 200 nm, less than 1% of the gland’s diameter. We recognize that flexure of the cuticle surface may result in variable crack width along its depth, restricting the true effective crack width [40, 41].

**Fig. 4.**
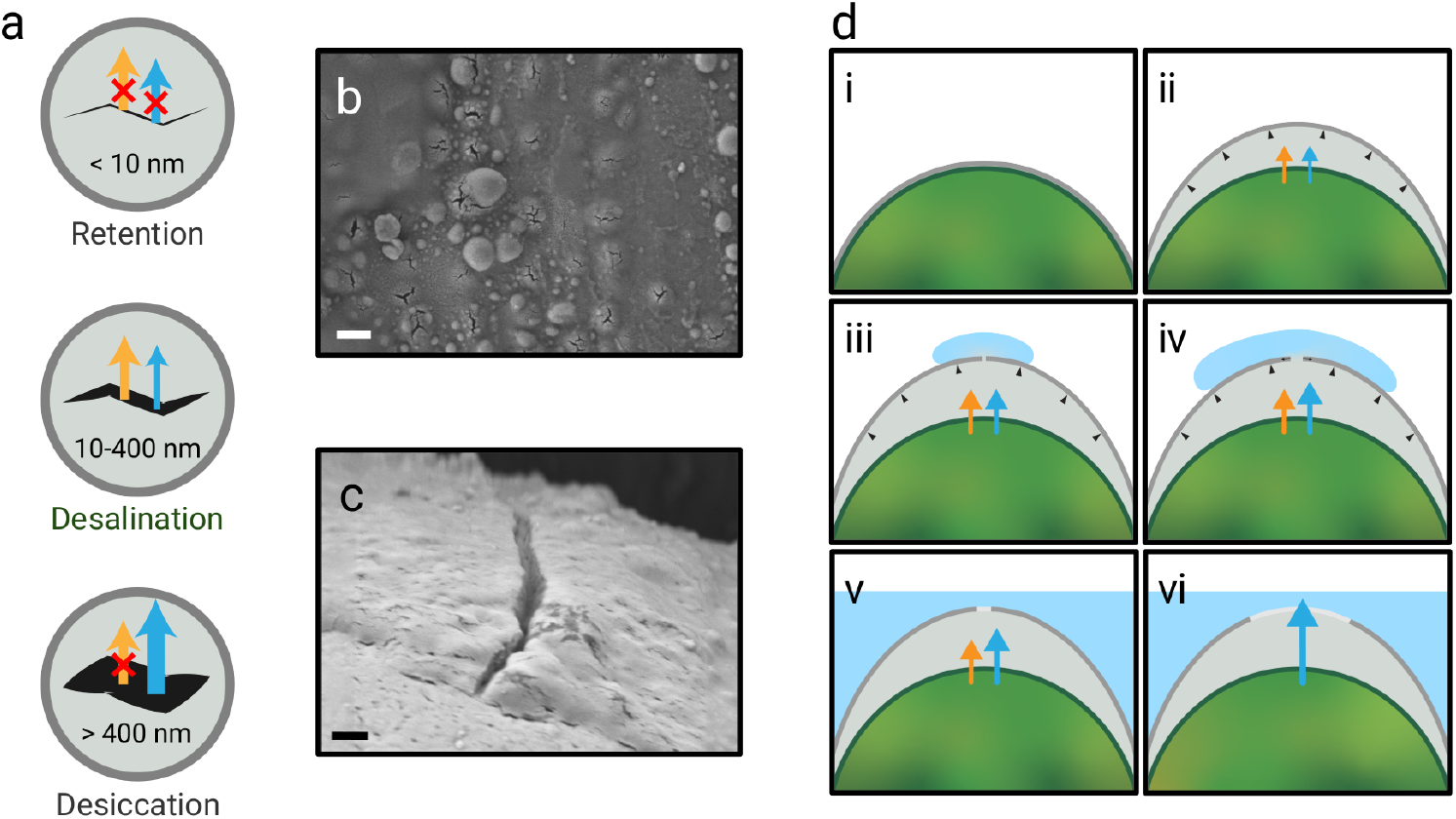
Fracture of the salt gland cuticle. a, Model predictions for functional regimes of cuticle crack sizes. Exceedingly narrow cracks (< 10 nm) restrict water flow and block salt efflux, leading to retention of salt and water. Intermediate cracks (c. 10-400 nm) allow both water and salt movement. Large cracks (> 400 nm) lead to impairment of ion transporters, arrest salt export while enabling catastrophic water loss. b-c, Cryo-SEM imaging reveals cracks in the cuticle surface through which brine escapes. cracks were observed on the order of 20 nm (b) or 100 nm (c) in width. d, Conceptual graphic of fracture in the cuticle. (i) The cuticle of the naive salt gland is crack-free, and the chamber has not yet fully inflated. (ii) As salt (orange) and water (blue) are exported into the subcuticular space, building pressure inflates the chamber. (iii) Fracture initiates when the pressure exceeds the cuticle strength. (iv) cracks grow until the chamber pressure drops below the propagation stress. (v) If cracks reach a stable size within the operable range of the salt gland, the leaf will be able to export salt without catastrophic water loss. (vi) If cracks grow too large, the salt gland exits the operable range, leading to desiccation. Scale bars: 500 nm (b,c)

Fig. 4d illustrates the development of cuticle fractures and the mechanics necessary to prevent excessive crack propagation and, ultimately, salt gland dysfunction. A naïve salt gland begins with an intact, crack-free cuticle surface. Upon activation of salt export, the flux of salt and water out of the cell inflates the chamber by pushing the flexible cuticle away from the rigid cell wall. When the chamber pressure exceeds the strength of the cuticle, a crack forms. The growing crack releases pressure in the chamber and eventually arrests. If the crack arrests within the operable range of the salt gland, salt can be exported without significant water loss. However, if the crack grows beyond the range of operable sizes, export efficiency will vanish as transporters become overwhelmed and osmotic movement of water from the gland leads to dysfunction and desiccation. This process is likely irreversible, as there are no known mechanisms for cuticle repair after its detachment from the cell wall. Moreover, we observed that leaves previously treated with salt continued to secrete fluid even after the watering regimen returned to exclusively deionized water. This implies a point of no return for salt gland function, which may contribute to the observed high leaf turnover rate of wild *N. mollis*.

How does the cuticle achieve a crack of intermediate size to stay within the desalinating range (Fig. 4a)? Exceedingly small cracks restrict water flow and block salt export, leading to retention of salt. Strength estimates for fruit cuticles suggest that the salt gland’s cuticle is not strong enough to resist the high pressures at the small crack limit [42, 43]. This apparent weakness is, in fact, essential for the crack to grow large enough to escape the retentive regime and activate salt efflux processes. To avoid runaway propagation into the desiccating regime, the crack’s growth must arrest within the intermediate range. The fractocohesive length is a length scale describing the dependence of a material’s vulnerability to crack propagation on the size of existing cracks. For a crack smaller than the fractocohesive length, the crack tip does not concentrate stress, and the strength of the material is independent of the crack’s size [44, 45]. We use the measured properties of fruit cuticles to estimate the fractocohesive length of the salt gland cuticle to be on the order of 1 *µ*m to 1 mm [44, 46, 47]. As this fractocohesive length is larger than the upper limit of the predicted operable range, and potentially larger than the length scale of the gland itself, we expect that the strength of the cuticle does not drop as the crack grows, allowing it to resist runaway propagation. Moreover, the pressure gradient across the cuticle vanishes with a crack size just above 100 nm, removing the driving force for propagation. Consequently, the crack arrests well within the operable range.

## DISCUSSION

Its persistent surface brine is a distinctive characteristic of *Nolana mollis*, but the risk of desiccation is not a unique problem. Even if the surface brine fully evaporates, the remaining surface salt content is not likely to be fully decoupled from the subcuticular space, due to its hygroscopic or deliquescent properties [48]. Here we use *N. mollis* as a model system because the concentration of its persistent brine can be directly measured, whereas the brine in contact with the glands cannot be directly sampled for plants like black mangroves (*Avicennia germinans*) whose leaf surfaces during the day are covered in salt but appear dry.

The limitations and design considerations explored here likely apply even to secreting glands without visibly detached cuticles [20]. Without a subcuticular chamber, the processes described in Fig. 2a would be confined to the hydrated cell wall. The inflated chamber can be considered simply as an extension of this space and provides a more convenient calculation of the water potential in which matric contributions from cell wall polymers can be ignored [21]. In non-chambered salt glands, the cuticle is often thickest around basal cells and can isolate secreting cells from adjacent epidermal cells, suggesting that hydraulic separation of the subcuticular space is important even in these systems [16].

There has been substantial interest in designing biomimetic desalination devices and engineering salt tolerance in crops, with significant effort dedicated to understanding the biological problems of regulating genes and engineering transporters for efficient salt transport [30, 49–52]. This work highlights that the physical problem of desalination must also be considered. Our model finds that structural solutions can circumvent energetic limitations on moving salt against a steep concentration gradient. Dilution of the intervening subcuticular chamber reduces the energetic cost of efflux and relies on the mechanical properties of the cuticle itself. Efforts to introduce similar strategies in other plants may benefit from a focus on structural properties instead of solely on transporter optimization.

Similar subcuticular chambers are found in oil-secreting glands, called elaiophores, in some floral structures [53–56] and as glandular trichomes, such as in tomato, among others [50, 57, 58]. Development of the cavities in these glands begins with digestion of the pectin strata to initially detach the cuticle and cell wall, followed by distention of the cuticle chamber with the accumulation of secreted oils [55]. It is likely that a similar process applies for the development of chambered salt glands, such as those recently described in some mangroves [21]. However, we note that elaiophores are designed for storage and mechanical disruption by either a pollinator or other visitor, and do not include unprovoked cuticle fracture as part of their development [59], as we propose is relevant for secreting salt glands. However, these structures may be potential candidates for endeavors seeking to establish desalination pathways in other plants.

We developed a model of the secreting salt gland to understand the interplay between the cuticle’s fracture mechanics and the biochemical processes involved in foliar desalination. Thermodynamic limitations to transporter energy balance suggest that ion transporters cannot be infinitely active. This constrains the functional parameter space of the salt gland, as both direct contact with the brine and highly restrictive flow across the cuticle lead to steep concentration gradients across the cell membrane and subsequent deactivation of export processes. Our model finds that desalination is achieved when cuticle crack size remains on the order of 10–400 nm, which we have confirmed through cryo-SEM imaging of functional glands. We estimate the cuticle’s fractocohesive length to be on the order of 1 *µ*m to 1 mm, well beyond the required range to avoid runaway crack propagation and remain within the operable region. Our work highlights the importance of the elasticity, strength, and fractocohesive length of the cuticle for secreting salt gland function and raises important questions regarding how plants achieve the appropriate values for these material properties to create a functional salt-exporting phenotype.

## MATERIALS AND METHODS

### Plant material

We use *Nolana mollis*, a shrub native to the Pan de Azúcar region of the Atacama Desert, Chile, which has been previously identified as a salt-secreting plant [25, 26]. Seeds were germinated according to the protocol presented by Ref. [60]. To germinate, seeds were first surface sterilized using a bleach (3% NaOCl), followed by 70% ethanol, then incubated in 500ppm gibberellic acid at room temperature for 5 to 7 days. After incubation, seeds were plated in groups of 15–30 onto Petri dishes lined with dampened circular filter paper. The plates were then covered with their clear lids and tilted at a small angle to prevent pooling of water around the seeds. A cotton ball soaked in water was also placed at the lower edge of the plate to maintain humidity. Plates were then loosely enclosed in clear plastic greenhouse trays and kept in a growth chamber with a 12h light/12h dark light cycle, 85% RH, and 21^°^C. Plates were monitored for three months. When a seed’s radicle reached 1–5cm long, the seedling was planted in a seedling cell with a mix of 60% potting mix (PRO-MIX BK, Premier Tech) and 40% coarse sand. Over the course of 12 months, seedlings were transferred to 4in and then 6in pots when necessary, and the relative humidity was reduced to 65% RH. All plants were watered exclusively with dH_2_O for at least one month prior to any treatments.

### Imaging of foliar surface features

Plants were bottom watered with 200 mL 100 mM NaCl 9 and 2 days prior to imaging to induce brine secretion. Branchlets were detached pre-dawn, wrapped loosely in damp paper towels, and kept in a plastic bag until they were processed for imaging at the Center for Nanoscale Systems (CNS) at Harvard University. Branchlets were directly imaged using a digital light microscope (VHX-7100, Keyence) with serial recording of 3D image stacks.

For cryo-SEM imaging, a leaf from the third to fifth whorl of the branch was extracted and rinsed with dH_2_O to remove surface salts and gently blotted dry with lint-free tissue paper (Kimwipe). The leaf was then submerged and frozen in degassed liquid nitrogen and mounted in a clamped sample holder with the adaxial surface exposed. Using a cryo-transfer shuttle (VCT 500, Leica), the sample was mounted in a Leica EM ACE600 to etch off surface ice for 10 min at -95^°^C. The sample was then cooled to -150^°^C and sputter coated with a 9 nm layer of platinum and palladium. The sample was transferred in the cryo-shuttle to be imaged at 3.0 kV in a FESEM (Zeiss Gemini 360) at -150^°^C.

### Confocal imaging of salt gland ultrastructure

Individual leaves of *N. mollis* were collected and fixed in 4% w/v acrolein (Polysciences, Warrington, PA, USA) in a modified piperazine-N, N’-bis (2-ethanesulfonic acid) (PIPES) buffer adjusted to pH 6.8 (50 mM PIPES and 1 mM MgSO_4_ from BDH, London, UK; and 5 mM EGTA) for 24 h. After rinsing them thrice in the buffer, 15 min each, the cellulose of the cell walls was stained with 0.01% w/v calcofluor white in 10 mM CHES buffer with 100 mM KCl (pH = 10) [61] for 1h, washed with water 10 min, and then counterstained with auramine O in M Tris/HCl buffer, (pH = 7.2), that labels cutinized lipids [62]. Then, the individual leaves were cleared with a solution containing ethanol:benzoyl benzoate 3:1 (v/v) for 6 h, ethanol:benzoyl benzoate 1:3 (v/v) and finally benzoyl benzoate:dibutyl phthalate 1:1 (v/v) for several days [63]. Whole mounts of leaves were carefully mounted onto slides with a small cavity, and imaged with a Zeiss LSM700 Confocal Microscope connected to an AxioCam 512 camera and Zen Blue 2.3 software, using a 63x/1.40 Oil DIC M27 Plan-Apochromat objective. 3D images were reconstructed from Z-stacks and assembled with the Image J1.51d software (National Institutes of Health, Bethesda, MD, USA).

### Leaf osmolality and water potential

Plants were divided into six groups: one control (dH_2_O) and five salt treatment groups (NaCl, KCl, LiCl, CaCl_2_, MgCl_2_, with electrical conductivity 16.45mS). Each group contained five plants and was split into two subgroups on opposite sides of the growth chamber to account for potential spatial variations in the growth chamber environment. Each plant was placed in a plastic saucer and bottom watered at dawn of day 0, with 200mL of either dH_2_O (Control) or its assigned salt solution. The saucers were covered with aluminum foil to limit evaporation before absorption. After day 2, all pots were flushed top-down with dH_2_O until the conductivity of the flowthrough was similar to that of the control.

Leaves were collected pre-dawn and placed into 0.5mL Eppendorf tubes pre-loaded with a filter paper disc to absorb the surface brine. These brine discs were then processed using a Wescor Vapro-5600 vapor pressure osmometer to determine osmolality of the surface brine.

To determine whole-leaf osmolality, leaves were washed with dH_2_O to remove surface salts, then gently blotted dry with lint-free tissue paper (Kimwipe). Leaves were snap frozen in liquid nitrogen and left to thaw for 15 minutes. Thawed leaves were again blotted dry to remove surface condensation, then loaded into an Eppendorf tube fitted with a 0.22-*µ*m pore cellulose acetate membrane spin filter (Costar 8161, Corning). Leaves were gently crushed with a spatula, careful not to disturb the filter. Samples were spun at 12,000 RPM for 10 minutes to extract liquid (Microfuge 18 and F241.5P Rotor, Beckman Coulter). 10 *µ*L of the extracted liquid was pipetted on a filter paper disc to determine osmolality with a vapor pressure osmometer (Wescor Vapro-5600).

On day 1 (24 h after treatment), additional leaves from three plants each from the control and NaCl groups were collected pre-dawn for whole-leaf water potential measurements. Leaves were rinsed with dH_2_O to remove surface salts and debris and blotted dry before loading 5 to 6 leaves into the chamber of a psychrometer. Samples were allowed to equilibrate for 16 to 24 hours in a water bath held at 25^°^C. Psychrometer measurements were obtained every 15 minutes with 10 seconds of cooling at -6000 *µ*amps using a CR6 datalogger (Campbell Scientific). The psychrometer voltage measurement was determined as the mean of the output plateau during evaporation. An exponential fit was calculated for each voltage time series to determine the equilibrated value, which was then converted to the corresponding water potential using the empirical calibration curves for each individual psychrometer.

### Mathematical model

We developed a steady-state mathematical model to describe the movement of water and salt through the chambered secreting salt gland. Model parameters are summarized in Table 1. The primary compartment of the model corresponds to the subcuticular chamber, with unknown concentration *c*_*c*_, pressure *P*_*c*_, and water potential *ψ*_*c*_. The water potential *ψ* = *P* − *cRT* describes the potential energy density of water. The chamber is connected to the gland’s secreting cell, with known water potential *ψ*_*g*_ and osmotic potential Π_*g*_ = *c*_*g*_ *RT* as determined above, and the surface brine, held at atmospheric pressure *P*_*b*_ = 0.1 MPa with a concentration *c*_*b*_, as determined above. We did not find significant differences in the leaf water potential or brine concentration across the various salt treatments, so we set these quantities at their mean observed values (Table 1).

**Table 1.** Model parameters.

| Parameter | Symbol | Value |
| --- | --- | --- |
| Ideal gas constant | $R$ | $8.314 \text{ J K}^{-1} \text{ mol}^{-1}$ |
| Temperature | $T$ | 298 K |
| Molar volume of water | $\bar{v}$ | $1.807 \times 10^{-5} \text{ m}^3 \text{ mol}^{-1}$ |
| Viscosity of water | $\eta$ | $10^{-3} \text{ Pa s}$ |
| Free diffusion of sodium | $D_{\text{free}}$ | $1.6 \times 10^{-9} \text{ m}^2 \text{ s}^{-1}$ [64] |
| Atmospheric pressure | $P_{\text{atm}}$ | 0.1 MPa |
| Gland water potential | $\psi_g$ | -1.5 MPa |
| Gland radius | $r$ | $15 \text{ } \mu\text{m}$ |
| Cuticle thickness | $l_c$ | $1 \text{ } \mu\text{m}$ [23] |
| Membrane permeability | $L_{gc}$ | $10^{-12} \text{ m}^2 \text{ s kg}^{-1}$ [65] |
| Brine concentration | $c_b$ | $2000 \text{ mmol kg}^{-1}$ |
| Transport gradient threshold | $\Delta c^*$ | $1400 \text{ mmol kg}^{-1}$ |
| Gland $\text{Na}^+$ concentration | $c_g$ | $100 \text{ mmol kg}^{-1}$ [3] |
| Maximum salt efflux | $J_t$ | $400 \text{ nmol m}^{-2} \text{ s}^{-1}$ [66] |
| Number of pores | $n$ | 1 |
| Activity steepness | $h$ | 100 |

Aquaporin-mediated movement of water between the secreting gland cell (g) and the chamber (c) is driven by the water potential gradient:

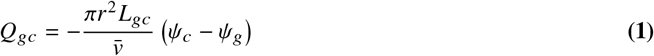

where *r* is the radius of the cell, *L*_*gc*_ is the aquaporin-mediated permeability to water of the cell wall and membrane, and 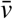 is the molar volume of water.

As the cuticle is not a selective membrane and is traversed by aqueous salt through cracks in the cuticle, we model the flux of water between the chamber and the brine (b) as a pressure-driven (advective) flux, where the brine is held at atmospheric pressure, *P*_atm_:

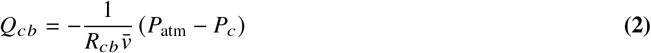

where *R*_*cb*_ is the resistance across the cuticle. We model the fractures as circular pores, so we describe this resistance through a perforated plate, following Ref. [67]:

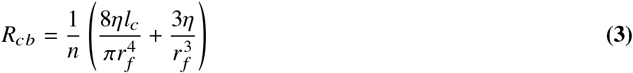

with *n* the number of pores, which for our primary analysis we take as 1, *η* the viscosity of water, *l*_*c*_ the thickness of the cuticle, and *r*_*f*_ the radius of the fracture.

Through these cracks, salt flux is coupled to water transport and described by an advection-diffusion equation:

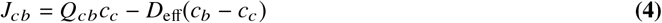

with the effective diffusion *D*_eff_ described as the diffusion through a perforated plate [68]:

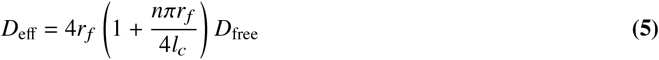

Active transporters mediate the secretion of salt from the living tissue into the chamber. While the precise nature of these processes is so far undetermined, we model this behavior as a function of the concentration gradient Δ*c*_*gc*_ = *c*_*c*_ − *c*_*g*_ between the chamber (*c*_*c*_) and gland (*c*_*g*_) and a threshold gradient value, Δ*c*^*^:

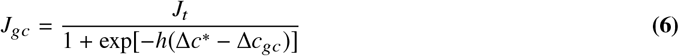

with *J*_*t*_ the maximum transport rate and *h* a coefficient to describe the steepness of the activation function. Other functions for the activity profile are explored in Fig. S2. We can estimate the activity threshold for ion transport through, as an example, a Na^+^/H^+^ antiporter [36] using the Nernst potential. The energy required to move one Na^+^ cation across the cell membrane is

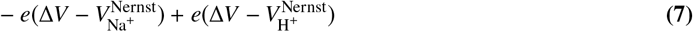

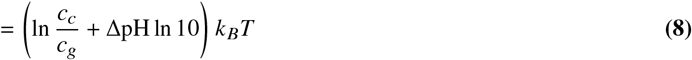

where Δ*V* is the membrane potential and *V* ^Nernst^ is the Nernst potential of each ion, *e* is the magnitude of the ion charge, and *k*_*B*_ is the Boltzmann constant. ATP provides around 19 *k*_*B*_*T* of energy to operate the transporter. Inducing the conformational changes for transporter function requires 8–16 *k*_*B*_*T* [33], limiting the remaining energy provided by ATP after the loss of some energy as heat. If the energy from Eq. 8 exceeds the remaining allotted energy, the transporter will not have enough energy to perform the transfer across the membrane. This required energy can be reduced by decreasing the pH of the chamber using separate proton pumps, which also require energetic input, or by reducing the concentration gradient. For our model, we estimate a concentration gradient of 1400 mM as the activity threshold, though we find that, as long as this threshold is above the lowest achievable gradient across the membrane, the gland is able to function (Fig. S2).

Sweeping across *r*_*f*_, which alters both *R*_*cb*_ and *D*_eff_, we use Newton’s method to solve for the steady state solution of the chamber’s concentration *c*_*c*_, pressure *P*_*c*_, and water potential *ψ*_*c*_, for which

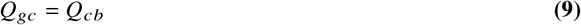

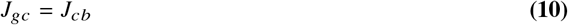

## ACKNOWLEDGMENTS

This work was supported by the National Science Foundation through the Harvard University Materials Research Science and Engineering Center (DMR-2011754). We acknowledge Adam Graham from the Harvard University Center for Nanoscale Systems (CNS) for his support on the cryo-SEM and light microsopy. MHM recognizes support from the Fannie and John Hertz Foundation Fellowship and the National Science Foundation Graduate Research Fellowship (Grant no. DGE1745303) and thanks Sophie Everbach, Liesbeth van den Brink, Tomás Fuenzalida, and Jacques Dumais for helpful discussions, and Ayman Fayad and Cory Hahn for greenhouse support. JML was funded by a grant from the Agencia Estatal de Investigación (PID2021-129074OB-100) and a Fulbright-CSIC fellowship (FULBR23036).

## REFERENCES

[1] A. R. Yeo and T. J. Flowers. Varietal differences in the toxicity of sodium ions in rice leaves. Physiologia Plantarum, 59:189–195, 1983. doi: 10.1111/j.1399-3054.1983.tb00756.x.

[2] Marilyn C. Ball and Rana Munns. Plant responses to salinity under elevated atmospheric concentrations of CO_2_. Australian Journal of Botany, 40(5): 515–525, 1992. doi: 10.1071/BT9920515.

[3] Timothy J. Flowers, Rana Munns, and Timothy D. Colmer. Sodium chloride toxicity and the cellular basis of salt tolerance in halophytes. Annals of Botany, 115(3):419–431, 2015. doi: 10.1093/aob/mcu217.

[4] Rana Munns, Richard A James, Matthew Gilliham, Timothy J Flowers, and Timothy D Colmer. Tissue tolerance: an essential but elusive trait for salt-tolerant crops. Functional Plant Biology, 43:1103–1113, 2016.

[5] Zohreh Heydarian, Min Yu, Margaret Gruber, Cathy Coutu, Stephen J. Robinson, and Dwayne D. Hegedus. Changes in gene expression in Camelina sativa roots and vegetative tissues in response to salinity stress. Scientific Reports, 8(1):1–22, 2018. doi: 10.1038/s41598-018-28204-4.

[6] Eva Van Zelm, Yanxia Zhang, and Christa Testerink. Salt tolerance mechanisms of plants. Annual Review of Plant Biology, 71(1):403–433, 2020. doi: 10.1146/annurev-arplant-050718.

[7] Paul M Hasegawa, Ray A Bressan, Jian-Kang Zhu, and Hans J Bohnert. Plant Cellular and Molecular Responses to High Salinity. Annual Review of Plant Physiology and Plant Molecular Biology, 51:463–499, 2000.

[8] Saverio Perri, Gabriel G. Katul, and Annalisa Molini. Xylem–phloem hydraulic coupling explains multiple osmoregulatory responses to salt stress. New Phytologist, 224(2):644–662, 2019. doi: 10.1111/nph.16072.

[9] Timothy J. Flowers, Hanaa K. Galal, and Lindell Bromham. Evolution of halophytes: multiple origins of salt tolerance in land plants. Functional Plant Biology, 37(2):604–612, 2010. doi: 10.1071/FP09269.

[10] John M. Cheeseman. The evolution of halophytes, glycophytes and crops, and its implications for food security under saline conditions. New Phytologist, 206(2):557–570, 2015. doi: 10.1111/nph.13217.

[11] J. Gorham. Photosynthesis, transpiration and salt fluxes through leaves of Leptochloa fusca L. Kunth. Plant, Cell & Environment, 10(2):191–196, 1987. doi: 10.1111/1365-3040.ep11602151.

[12] Marilyn C. Ball. Salinity tolerance in the mangroves Aegiceras corniculatum and Avicennia marina. I. Water use in relation to growth, carbon partitioning, and salt balance. Australian Journal of Plant Physiology, 15(3):447–464, 1988. doi: 10.1071/PP9880447.

[13] Timothy J. Flowers and Timothy D. Colmer. Salinity tolerance in halophytes. New Phytologist, 179(4):945–963, 2008. doi: 10.1111/j.1469-8137.2008.02531.x.

[14] Mingjing Zhang, Yanlu Liu, Guoliang Han, Yi Zhang, Baoshan Wang, and Min Chen. Salt tolerance mechanisms in trees: research progress. Trees, 35(3):717–730, 2021. doi: 10.1007/s00468-020-02060-0.

[15] P. B. Tomlinson. Water relations and salt balance. In The botany of mangroves, pages 116–130. Cambridge University Press, 1986. ISBN 9780521255677.

[16] Maheshi Dassanayake and John C. Larkin. Making plants break a sweat: The structure, function, and evolution of plant salt glands. Frontiers in Plant Science, 8(406):1–20, 2017. doi: 10.3389/fpls.2017.00406.

[17] Siegmar-W. Breckle. Salinity tolerance of different halophyte types. In N. El Bassam, M. Dambroth, and B.C. Loughman, editors, Genetic Aspects of Plant Mineral Nutrition, pages 167–175. Springer, Dordrecht, 1990. doi: 10.1007/978-94-009-2053-8_26.

[18] Feng Ding, Jian-Chao Yang, Fang Yuan, and Bao-Shan Wang. Progress in mechanism of salt excretion in recretohalopytes. Frontiers in Biology, 5(2): 164–170, 2010. doi: 10.1007/s11515-010-0032-7.

[19] N Campbell and W W Thomson. The Ultrastructure of Frankenia Salt Glands. Annals of Botany, 40(4):681–686, 1976. doi: 10.1093/oxfordjournals.aob.a085181.

[20] Fang Yuan, Bingying Leng, and Baoshan Wang. Progress in Studying Salt Secretion from the Salt Glands in Recretohalophytes: How Do Plants Secrete Salt? Frontiers in Plant Science, 7(977):1–12, 2016. doi: 10.3389/fpls.2016.00977.

[21] Bing-Jie Chi, Ze-Jun Guo, Ming-Yue Wei, Shi-Wei Song, You-Hui Zhong, Jing-Wen Liu, Yu-Chen Zhang, Jing Li, Chao-Qun Xu, Xue-Yi Zhu, and Hai-Lei Zheng. Structural, developmental and functional analyses of leaf salt glands of mangrove recretohalophyte Aegiceras corniculatum. Tree Physiology, 44(1):tpad123, 2023. doi: 10.1093/treephys/tpad123.

[22] Yunquan Deng, Zhongtao Feng, Fang Yuan, Jianrong Guo, Shanshan Suo, and Baoshan Wang. Identification and functional analysis of the autofluorescent substance in Limonium bicolor salt glands. Plant Physiology and Biochemistry, 97:20–27, 2015. doi: 10.1016/j.plaphy.2015.09.007.

[23] C. Alice Levering and William W. Thomson. The ultrastructure of the salt gland of Spartina foliosa. Planta, 97(3):183–196, 1971. doi: 10.1007/BF00389200.

[24] Zouhaier Barhoumi, Wahbi Djebali, Chedly Abdelly, Wided Chaïbi, and Abderrazak Smaoui. Ultrastructure of Aeluropus littoralis leaf salt glands under NaCl stress. Protoplasma, 233(3-4):195–202, 2008. doi: 10.1007/s00709-008-0003-x.

[25] H. A. Mooney, S. L. Gulmon, J. Ehleringer, and P. W. Rundel. Atmospheric water uptake by an Atacama Desert shrub. Science, 209(4457):693–694, 1980. doi: 10.1126/science.209.4457.693.

[26] P. W. Rundel, James R. Ehleringer, H. A. Mooney, and S. L. Gulmon. Patterns of Drought Response in Leaf-Succulent Shrubs of the Coastal Atacama Desert in Northern Chile. Oecologia, 46:196–200, 1980.

[27] Jess Gersony, Anju Manandhar, Uri Hochberg, Nora Abdellaoui, Paula Llanos, Jacques Dumais, N Michele Holbrook, and Fulton E Rockwell. Making dew in the Atacama Desert: a paradigmatic case of plant water uptake water from an unsaturated atmosphere fails a test. Annals of Botany, 2024. doi: 10.1093/aob/mcae221.

[28] Rainer Hedrich. Ion channels in plants. Physiological Reviews, 92(4):1777–1811, 2012. doi: 10.1152/physrev.00038.2011.

[29] Bernhard Huchzermeyer and Tim Flowers. Putting halophytes to work-genetics, biochemistry and physiology. Functional Plant Biology, 40(9), 2013. doi: 10.1071/FPv40n9-FO.

[30] Jesper T. Pedersen and Michael Palmgren. Why do plants lack sodium pumps and would they benefit from having one? Functional Plant Biology, 44 (5):473–479, 2017. doi: 10.1071/FP16422.

[31] Timothy J. Flowers, Edward P. Glenn, and Vadim Volkov. Could vesicular transport of Na^+^ and Cl^−^ be a feature of salt tolerance in halophytes? Annals of Botany, 123(1):1–18, 2019. doi: 10.1093/aob/mcy164.

[32] Michael R. Blatt. A charged existence: A century of transmembrane ion transport in plants. Plant Physiology, 195(1):79–110, 2024. doi: 10.1093/plphys/kiad630.

[33] Fabrizio Marinelli and José D. Faraldo-Gómez. Conformational free-energy landscapes of a Na^+^/Ca^2+^ exchanger explain its alternating-access mechanism and functional specificity. Proceedings of the National Academy of Sciences, 121(16):e2318009121, 2024. doi: 10.1073/pnas.2318009121.

[34] Alessio Accardi, Michael Walden, Wang Nguitragool, Hariharan Jayaram, Carole Williams, and Christopher Miller. Separate ion pathways in a Cl^−^/H^+^ exchanger. Journal of General Physiology, 126(6):563–570, 2005. doi: 10.1085/jgp.200509417.

[35] Qing Xie, Yang Zhou, and Xingyu Jiang. Structure, Function, and Regulation of the Plasma Membrane Na^+^/H^+^ Antiporter Salt Overly Sensitive 1 in Plants. Frontiers in Plant Science, 13(March), 2022. doi: 10.3389/fpls.2022.866265.

[36] Xiang-yun Zhang, Ling-hui Tang, Jia-wei Nie, Chun-rui Zhang, Xiaonan Han, Qi-yu Li, Li Qin, Mei-hua Wang, Xiahe Huang, Feifei Yu, Min Su, Yingchun Wang, Rui-ming Xu, Yan Guo, Qi Xie, and Yu-hang Chen. Structure and activation mechanism of the rice Salt Overly Sensitive 1 (SOS1) Na^+^/H^+^ antiporter. Nature Plants, 9(11):1924–1936, 2023. doi: 10.1038/s41477-023-01551-5.

[37] W. W. Thomson. The Structure and Function of Salt Glands. In Plants in Saline Environments, pages 118–146. Springer-Verlag, Berlin, 1975. doi: 10.1007/978-3-642-80929-3_9.

[38] Masahiro Koyama and Takao Oi. Morphology and excreting-function of microhairs in salt-tolerant Zoysia japonica, comparing adaxial and abaxial leaf surfaces. Flora, 312:152472, 2024. doi: 10.1016/j.flora.2024.152472.

[39] Richard G Allen, Luis S Pereira, Dirk Raes, Martin Smith, and Others. Crop evapotranspiration-Guidelines for computing crop water requirements-FAO Irrigation and drainage paper 56. FAO, 300(9):D05109, 1998.

[40] Anneline H. Christensen and Kaare H. Jensen. Viscous flow in a slit between two elastic plates. Physical Review Fluids, 5(4):1–10, 2020. doi: 10.1103/PhysRevFluids.5.044101.

[41] Jean François Louf, Jan Knoblauch, and Kaare H. Jensen. Bending and stretching of soft pores enable passive control of fluid flows. Physical Review Letters, 125(9):98101, 2020. doi: 10.1103/PhysRevLett.125.098101.

[42] Hendrik Bargel and Christoph Neinhuis. Tomato (Lycopersicon esculentum Mill.) fruit growth and ripening as related to the biomechanical properties of fruit skin and isolated cuticle. Journal of Experimental Botany, 56(413):1049–1060, 2005. doi: 10.1093/jxb/eri098.

[43] Eva Dominguez, José Alejandro Heredia-Guerrero, and Antonio Heredia. The biophysical design of plant cuticles: An overview. New Phytologist, 189(4):938–949, 2011. doi: 10.1111/j.1469-8137.2010.03553.x.

[44] Chao Chen, Zhengjin Wang, and Zhigang Suo. Flaw sensitivity of highly stretchable materials. Extreme Mechanics Letters, 10:50–57, 2017. doi: 10.1016/j.eml.2016.10.002.

[45] Jie Ma, Xizhe Zhang, Daochen Yin, Yijie Cai, Zihang Shen, Zhi Sheng, Jiabao Bai, Shaoxing Qu, Shuze Zhu, and Zheng Jia. Designing Ultratough Single-Network Hydrogels with Centimeter-Scale Fractocohesive Lengths via Inelastic Crack Blunting. Advanced Materials, 36(23):1–11, 2024. doi: 10.1002/adma.202311795.

[46] Antonio J. Matas, Eward D. Cobb, Dominick J. Paolillo, and Karl J. Niklas. Crack resistance in cherry tomato fruit correlates with cuticular membrane thickness. HortScience, 39(6):1354–1358, 2004.

[47] Laura España, José A. Heredia-Guerrero, Patricia Segado, José J. Benítez, Antonio Heredia, and Eva Domínguez. Biomechanical properties of the tomato (Solanum lycopersicum) fruit cuticle during development are modulated by changes in the relative amounts of its components. New Phytologist, 202(3):790–802, 2014. doi: 10.1111/nph.12727.

[48] Huanhuan Zhang, Wenjun Gu, Yong Jie Li, and Mingjin Tang. Hygroscopic properties of sodium and potassium salts as related to saline mineral dusts and sea salt aerosols. Journal of Environmental Sciences, 95:65–72, 2020. doi: 10.1016/j.jes.2020.03.046.

[49] Timothy John Flowers, Aurora Garcia, Mikiko Koyama, and Anthony Richard Yeo. Breeding for salt tolerance in crop plants - the role of molecular biology. Acta Physiologiae Plantarum, 19(4):427–433, 1997.

[50] Joris J. Glas, Bernardus C.J. Schimmel, Juan M. Alba, Rocío Escobar-Bravo, Robert C. Schuurink, and Merijn R. Kant. Plant glandular trichomes as targets for breeding or engineering of resistance to herbivores. International Journal of Molecular Sciences, 13(12):17077–17103, 2012. doi: 10.3390/ijms131217077.

[51] Lukasz Kotula, Pedro Garcia Caparros, Christian Zörb, Timothy D. Colmer, and Timothy J. Flowers. Improving crop salt tolerance using transgenic approaches: An update and physiological analysis. Plant Cell and Environment, 43(12):2932–2956, 2020. doi: 10.1111/pce.13865.

[52] Maria Di Vincenzo, Alberto Tiraferri, Valentina Elena Musteata, Stefan Chisca, Rachid Sougrat, Li Bo Huang, Suzana P. Nunes, and Mihail Barboiu. Biomimetic artificial water channel membranes for enhanced desalination. Nature Nanotechnology, 16(2):190–196, 2021. doi: 10.1038/s41565-020-00796-x.

[53] Beryl B Simpson, John L Neff, and David S Seigler. Floral Biology and Floral Rewards of Lysimachia (Primulaceae). The American Midland Naturalist, 110(2):249–256, 1983.

[54] Malgorzata Stpiczyńska and Kevin L. Davies. Elaiophore structure and oil secretion in flowers of Oncidium trulliferum Lindl. and Ornithophora radicans (Rchb.f.) Garay Pabst (Oncidiinae: Orchidaceae). Annals of Botany, 101(3):375–384, 2008. doi: 10.1093/aob/mcm297.

[55] Clivia Carolina Fiorilo Possobom, Elza Guimarães, and Silvia Rodrigues Machado. Structure and secretion mechanisms of floral glands in Diplopterys pubipetala (Malpighiaceae), a neotropical species. Flora, 211:26–39, 2015.

[56] Clivia Carolina Fiorilo Possobom and Silvia Rodrigues Machado. Elaiophores: Their taxonomic distribution, morphology and functions. Acta Botanica Brasilica, 31(3):503–524, 2017. doi: 10.1590/0102-33062017abb0088.

[57] Ryan W. Shepherd and George J. Wagner. Phylloplane proteins: emerging defenses at the aerial frontline? Trends in Plant Science, 12(2):51–56, 2007. doi: 10.1016/j.tplants.2006.12.003.

[58] Robert Schuurink and Alain Tissier. Glandular trichomes: micro-organs with model status? New Phytologist, 225(6):2251–2266, 2020. doi: 10.1111/nph.16283.

[59] A.N Sérsic. Pollination Biology in the Genus Calceolaria L.(Calceolariaceae). Stapfia, 82:1–121, 2004.

[60] Liesbeth van den Brink, Melissa H Mai, Lorenz Henneberg, Margret Ecke, and Rafaella Canessa. Gibberellin enhances germination of nolana mollis and heliotropium pycnophyllum seeds. bioRxiv, 2025. doi: 10.1101/2025.01.10.632421.

[61] J. Hughes and Margaret E. McCully. The Use of an Optical Brightener in the Study of Plant Structure. Stain Technology, 50(5):319–329, 1975. doi: 10.3109/10520297509117082.

[62] Yolande Heslop-Harrison and K. R. Shivanna. The receptive surface of the angiosperm stigma. Annals of Botany, 41(6):1233–1258, 11 1977. ISSN 0305-7364. doi: 10.1093/oxfordjournals.aob.a085414.

[63] Charles F. Crane and John G. Carman. Mechanisms of Apomixis in Elymus rectisetus from Eastern Australia and New Zealand. American Journal of Botany, 74(4):477–496, 1987. doi: 10.2307/2443827.

[64] E. A. Guggenheim. The diffusion coefficient of sodium chloride. Transactions of the Faraday Society, 50:1048–1051, 1954. doi: 10.1039/tf9545001048.

[65] Paul J. Kramer and John S. Boyer. Water Relations of Plants and Soils. Academic Press, Inc., New York, 1 edition, 1995. ISBN 9780124250604. doi: 10.1097/00010694-199604000-00007.

[66] John W. Oross, Robert T. Leonard, and William W. Thomson. Flux rate and a secretion model for salt glands of grasses. Israel Journal of Botany, 34 (2-4):69–77, 1985. doi: 10.1080/0021213X.1985.10677012.

[67] Kaare Hartvig Jensen, Daniel Leroy Mullendore, Noel Michele Holbrook, Tomas Bohr, Michael Knoblauch, and Henrik Bruus. Modeling the hydrodynamics of phloem sieve plates. Frontiers in Plant Science, 3(3):1–11, 2012. doi: 10.3389/fpls.2012.00151.

[68] William F Pickard. Lumped Circuit Approximations for Flux in Systems Governed by the Laplace Equation. Chemical Engineering Science, 36: 191–197, 1980.

