## Supplemental Information for "Secreting salt glands constrain cuticle fracture to enhance desalination efficiency"

### SUPPLEMENTARY MATERIALS FOR
### SECRETING SALT GLANDS CONSTRAIN CUTICLE FRACTURE TO ENHANCE DESALINATION EFFICIENCY

**Authors:** Melissa H. Mai, Fulton E. Rockwell, Juan M. Losada, Nya Nicholson, Zhigang Suo, N. Michele Holbrook

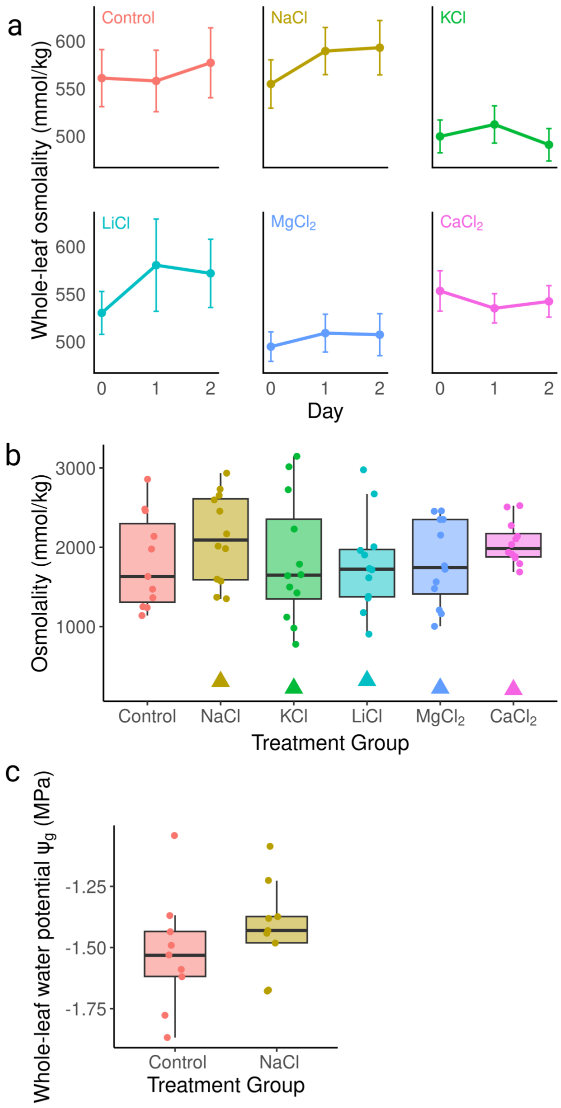


**Fig. S1.** Leaf osmolality and water potential for *Nolana mollis* following salt treatments applied on day 0. a, Whole-leaf osmolality measurements show no change after treatment across all group. Data were collected for 5 plants per group over 3 trials. Error bars denote SE. b, Pre-dawn surface brine concentration (boxes, round points) on day 2. Triangles refer to the measured osmolality of the treatment solutions (c. 300 mmol/kg). Due to greenhouse facility conditions, all plants were watered for several weeks with tap water, leading to brine secretion in the control group. At least one month prior to treatment, all plants were watered exclusively with deionized water, but leaves on plants in the control group continued to secrete brine, though at qualitatively smaller volumes than those in the salt treatment groups. c, Whole-leaf water potential $\psi_g$ for leaves in the control and NaCl groups on day 1.

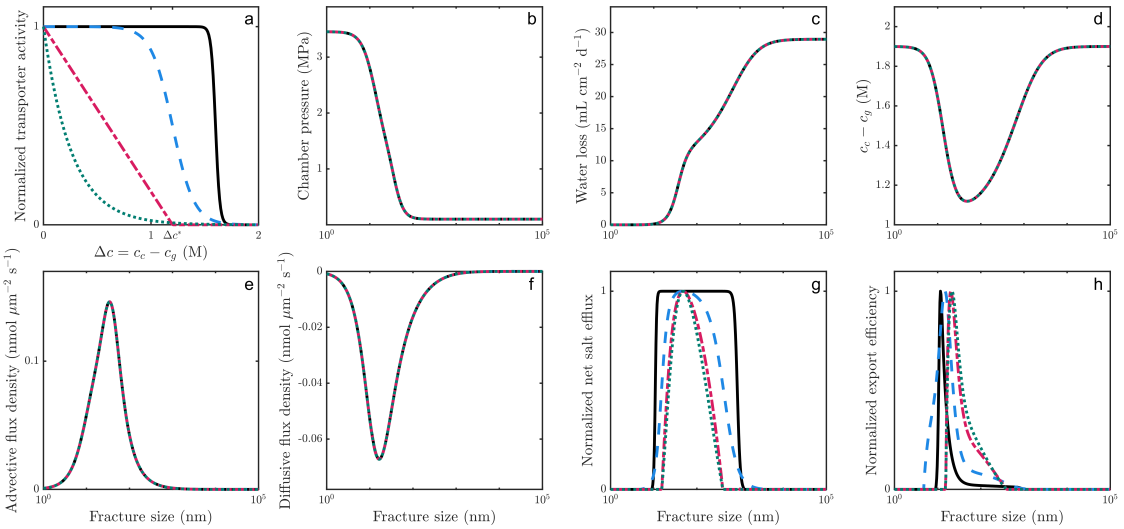


**Fig. S2.** Effect of various transporter activity profiles on chambered salt gland function. The solid black lines corresponds to a profile with a higher gradient threshold. a, Transporter activity as a function of the transmembrane concentration gradient. b, Chamber pressure. c, Total gland water loss. d, Transmembrane concentration gradient. e-f, Advective (e) and diffusive (f) flux densities, normalized by crack area. g, Normalized salt efflux rates. h, Normalized export efficiency.

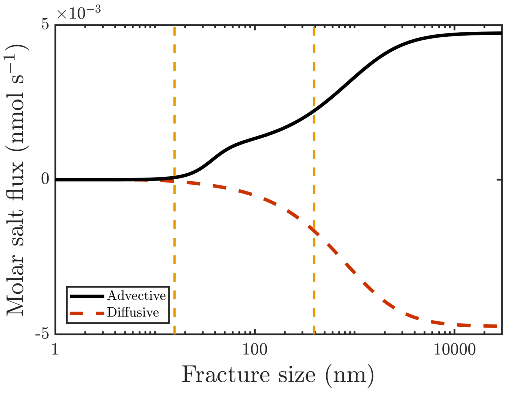


**Fig. S3.** Total advective (solid black) and diffusive (dashed red) fluxes across the cuticle as a function of crack size. The area between the vertical dashed lines correspond to the crack sizes that allow for positive salt export.
